# Identification of PS1/gamma-secretase and glutamate transporter GLT-1 interaction sites

**DOI:** 10.1101/2023.05.30.542955

**Authors:** Florian Perrin, Priyanka Sinha, Shane Mitchell, Masato Maesako, Oksana Berezovska

## Abstract

The recently discovered interaction between Presenilin 1 (PS1), a catalytic subunit of γ-secretase responsible for generating amyloid-β (Aβ) peptides, and GLT-1, a major glutamate transporter in the brain (EAAT2) provides a mechanistic link between these two key factors involved in Alzheimer’s disease (AD) pathology. Modulating this interaction can be crucial to understand the consequence of such crosstalk in AD context and beyond. However, the interaction sites between these two proteins are unknown. Herein, we utilized an alanine scanning approach coupled with FRET-based fluorescence lifetime imaging microscopy (FLIM) to identify the interaction sites between PS1 and GLT-1 in their native environment within intact cells. We found that GLT-1 residues at position 276 to 279 (TM5) and PS1 residues at position 249 to 252 (TM6) are crucial for GLT-1/PS1 interaction. These results have been cross validated using AlphaFold Multimer prediction. To further investigate whether this interaction of endogenously expressed GLT-1 and PS1 can be prevented in primary neurons, we designed PS1/GLT-1 cell-permeable peptides (CPPs) targeting the PS1 or GLT-1 binding site. We used HIV TAT domain to allow for cell penetration which was assayed in neurons. First, we assessed the toxicity and penetration of CPPs by confocal microscopy. Next, to ensure the efficiency of CPPs, we monitored the modulation of GLT-1/PS1 interaction in intact neurons by FLIM. We saw significantly less interaction between PS1 and GLT-1 with both CPPs. Our study establishes a new tool to study the functional aspect of GLT-1/PS1 interaction and its relevance in normal physiology and AD models.

## Introduction

One of the hallmarks of the neurodegenerative disease Alzheimer’s disease (AD) is the formation of extracellular amyloid-beta (Aβ) plaques. Presenilin 1 (PS1), a catalytic subunit of γ-secretase that cleaves amyloid precursor protein (APP) to produce Aβ peptides, is an established critical player in developing AD pathology (1, 2). Several studies have linked PS1/amyloid pathology to glutamate activity (3-5), although the precise molecular and mechanistic link is unclear. Hyperactivation and impaired deactivation of the default-mode networks with deficiencies in glutamate transport occur at early stages of AD (6-8); these features correlate with the loss of synaptic function and cognitive decline in AD patients (9, 10). Patients with childhood epilepsy, reexamined 50 years later, presented an increase in amyloid load, a risk factor for developing AD later in life (11). Furthermore, AD mouse model studies show that the neuronal network hyperactivity occurs prior to Aβ deposition, suggesting that glutamate malfunction is one of the earliest dysfunctions in the Alzheimer’s pathological cascade (12, 13). Conversely, AD patients, particularly those with mutations in PS1, have a higher incidence of epileptiform activity (6, 14-17). The occurrence of seizures is increased among cognitively asymptomatic fAD mutation carriers (18).

Interestingly, while searching for potential interactors of PS1, we identified a novel interaction between PS1 and glutamate transporter 1 (GLT-1 a.k.a. excitatory amino acid transporter 2, EAAT2) (19), a major glutamate transporter in the brain (20, 21). We reported that GLT-1 directly binds to PS1/γ-secretase (19), providing a possible mechanistic link between the PS1/amyloid pathology and GLT-1/hyperactivity in the brain, and solidifying the hypothesis that GLT-1 is involved in neuronal hyperexcitability in AD. We propose that the PS1/GLT-1 interaction is functional and may become dysregulated in AD (17, 19, 22). However, to test this hypothesis and to establish the significance of this interaction, we first need to discover the precise interaction site(s) between GLT-1 and PS1.

Thus, this study aimed to identify the exact interaction sites between PS1 and GLT-1 using an alanine scanning strategy in cell lines. A fluorescence lifetime imaging microscopy (FLIM) approach was used to evaluate the interaction between PS1 and GLT-1 in intact cells expressing either wild type or alanine mutant PS1 and GLT-1. Two potential sites on GLT-1 and one on PS1 that comprise four amino acids each have been identified; alanine mutations in these sites resulted in significantly decreased interaction between PS1 and GLT-1. These results are also in accord with the predicted AlphaFold multimers(23, 24), which allowed us to establish a model of human GLT-1/PS1 interaction.

Based on the sequence of these potential interaction sites, we designed cell permeable peptides (CPPs) by using the TAT domain from HIV to allow small peptides to penetrate cells. Inhibition of this interaction by CPPs was assessed in primary neurons which express PS1 and GLT-1 endogenously. We found that the interaction is disrupted with both CPPs targeting either PS1 or GLT-1. These CPPs can be used as a tool to modulate GLT-1/PS1 interaction to study its functional impact.

## Materials and Methods

### Cell cultures and transfection

Chinese hamster ovary (CHO) cells were maintained in OPTI MEM medium supplemented with 5% FBS in a 37°C CO_2_ incubator. Human embryonic kidney (HEK) cells in which PS1 and PS2 are knocked down (HEK PS DKO) were kindly provided by Dr. Dennis Selkoe, BWH, Boston, MA, and were maintained in DMEM supplemented with 5% FBS and 1% GlutaMax (Life Technologies, Carlsbad, CA) in a 37°C CO_2_ incubator. Lipofectamine 3000 (Life Technologies, Carlsbad, CA) was used for transient transfection according to the manufacturer’s instructions. Mixed cortical primary neurons from 16 to 18 embryonic-day-old embryos were enzymatically dissociated using the papain dissociation system (Worthington Biochemical Corporation, Lakewood, NJ). The neuronal cultures were maintained in Neurobasal medium supplemented with 2% B27 supplement, 1% GlutaMax, and 1% penicillin/streptomycin (ThermoScientific, Waltham, MA) in a 37 °C, 5% CO_2_ incubator. Primary neurons were maintained in culture and treated at 10–12 days in vitro (DIV).

### Plasmid constructs

A plasmid encoding PS1wt was cloned into pcDNA3.1 vector (Addgene). The GLT-1 encoding sequence was subcloned into pcDNA™6 V5 Myc (Thermo Scientific).

Alanine scanning on PS1wt and GLT-1wt was performed by cloning with primers provided by Twist Bioscience.

### Chemicals and treatments

PS1/GLT-1 interaction was blocked by incubating primary neurons for 2 h at 37 °C with 5 μM CPPs. The CPP containing the GLT-1 sequence involved in binding to PS1 is referred to as GLT-1 CPP; the CPP with PS1 sequence implicated in interaction with GLT-1 is referred to as PS1 CPP. GLT-1 CPP was obtained by fusing amino acids (aa) 47–57 from the HIV1 TAT protein (YGRKKRRQRRR) with the PS1 peptide WLILAVIS through a GGG linker. The GLT-1 CPP was obtained by fusing the same HIV1 TAT peptide (YGRKKRRQRRR) with the GLT-1-derived sequence NEIVMKLV through a GGG linker. The same HIV1 TAT peptide fused through a GGG linker to a scramble sequence MKLVIVEN or IWSLLAIV for PS1 and GLT-1 respectively, were used as negative controls. The peptides were synthesized at the MGH peptide core facility https://researchcores.partners.org/pepcor/about.

### Immunocytochemistry (ICC)

For ICC, *in vitro* cultured cells were washed twice with Dulbecco’s Phosphate Buffered Saline without Calcium Chloride or Magnesium Chloride (PBS; Thermo Scientific, Waltham, MA) and fixed by 10-min incubation with 4% PFA. Following the fixation, cells were blocked with 1% normal donkey serum (NDS; Jackson ImmunoResearch labs, West Grove, PA). After three 5 min washes in PBS, samples were incubated overnight with respective primary antibodies (1:200) in 1.5% NDS. Excess primary antibodies were washed off with three 5 min washes in PBS, and the corresponding Alexa Fluor 488- or Cy3-conjugated secondary antibodies (1:500) were applied for an hour at room temperature. Cells and tissue were then washed three additional times in PBS. Cells were coverslipped with VectaShield mounting medium (Vector Laboratories, Inc., Burlingame, CA).

### Antibodies

The following primary antibodies were used: guinea pig anti-GLT-1 (AB1783, EMD Millipore, Temecula, CA) for ICC, rabbit anti-GLT-1 (ab41621, Abcam, Cambridge, MA) for Western blot, rabbit anti-PS1 raised against aa 263–378 of PS1 (S182, Sigma-Aldrich, St. Louis, MO); mouse anti-β-actin (A2228, Sigma-Aldrich, St. Louis, MO). Alexa Fluor 488 (Thermo Scientific, Waltham, MA) and Cy3-conjugated secondary antibodies (Jackson ImmunoResearch, West Grove, PA) were applied for confocal microscopy imaging, and IRDye680/800-labelled ones (Li-COR, Lincoln, NE) were used for western blotting.

### Cytotoxicity assay

Cytotoxicity was analyzed using the lactate dehydrogenase (LDH) cytotoxicity assay (Roche, Indianapolis, IN). Briefly, conditioned medium was collected from the respective wells, mixed with the assay solution, incubated for 20 min in the dark, and the absorbance at 490 nm was measured using a spectrophotometer. For a positive control, cells were incubated for 45 min at 37 °C with 1% Triton X (TX)-100.

### Fluorescence lifetime imaging microscopy (FLIM)

The FLIM assay was conducted as described previously (25). Briefly, cells were immunostained with anti-GLT-1 and anti-PS1 C-terminal antibodies. Corresponding secondary antibodies conjugated with Alexa Fluor 488 (AF488) and Cy3 fluorophores were used as the donor and acceptor, respectively. The sample where the acceptor antibody was omitted was used as a negative control to record the baseline lifetime (*t*1) of the donor fluorophore. A femtosecond-pulsed Spectra-Physics Mai Tai laser (Spectra-Physics, Milpitas, CA) at 850 nm was used for two-photon fluorescence excitation. AF488 fluorescence was acquired using an emission filter centered at 515/30 nm. The donor fluorophore lifetimes were measured with a high-speed photomultiplier tube (MCP R3809; Hamamatsu, Bridgewater, NJ) and a fast time-correlated single-photon counting acquisition board (SPC-830; Becker & Hickl, Berlin, Germany). The images were analyzed using SPCImage software (Becker and Hickl, Berlin, Germany). The AF488 lifetimes were calculated by fitting raw data to the single-exponential (AF488 negative control) or multi-exponential (AF488- and Cy3-double immunostained sample) fluorescence decay curves. The *t*_*2*_ lifetime values that are shorter than *t*_*1*_ indicate the presence of FRET, i.e., less than 5–10 nm distance between the donor and acceptor fluorophores. FRET efficiency was calculated by subtracting the measured lifetime (*t2*) from the baseline lifetime (*t1*), divided by *t*1, expressed as a percentage (i.e. [(*t1*-*t2*)/*t1*]*100).

### Western blotting

The samples were loaded on 4–12% Bis-Tris NuPage polyacrylamide gels (Thermo Fisher Scientific, Waltham, MA) and transferred to an iBlot™ 2 Transfer Stack nitrocellulose (Thermo Fisher Scientific) iBlot™ 2 Gel Transfer Device. The detection was performed by immunoblotting with specific primary (1:1000) and corresponding IRdye680/800-conjugated secondary antibodies (1:5000), and the bands were visualized using an Odyssey Infrared Imaging System (Li-COR, Lincoln, NE). The quantitative analysis of the respective bands’ optical density was performed using ImageStudio Lite Ver 5.2 software.

### Protein 3D models

GLT-1 and PS1 sequences were profiled for secondary structure, intrinsic disorder, and accessibility propensities with AlphaFold2 and AlphaFold multimer. Closest templates were retrieved based on the top 5 pLDDT ranking. Templates from AlphaFold2 were used to effectively build the 3D models of the interaction interface between PS1 and GLT-1. Images were prepared in PyMOL Molecular Graphics System.

### Statistical Analysis

Statistical analysis was performed using GraphPad Prism 9 software (La Jolla, CA). ANOVA with Bonferroni’s post-hoc correction was used unless otherwise stated in the results. Values were considered significant at *P*<0.05.

## Results

### Expression of PS1 and GLT-1 alanine mutants

To efficiently identify potential interaction sites between PS1 and GLT-1, we performed an alanine scan on several windows of residues taking cues from the juxta-transmembrane region of PS1 and GLT-1 as both are polytopic membrane proteins. Our strategy was to mutate three to four amino acids (aa) to alanine at a time, based on the hydrophobicity and accessibility of folded PS1 and GLT-1. Mutated amino acids are shown in bold with each color corresponding to a set of residues mutated as a group while unmutated residues are in standard font in light green (Figure 1A). Expression of the GLT-1 and PS1 mutants has been assayed by western blot by transient transfection in HEK PS DKO cells (Figure 1B). To analyze the expression of GLT-1 or PS1 alanine mutants, we performed co-transfection of the mutants with pcDNA and PS1 wt and GLT-1 wt, respectively. Western blotting was performed to verify that the mutants are indeed expressing and do not modify PS1wt or GLT-1wt expression. Membranes were probed with GLT-1, PS1, and β-actin antibodies. All PS1 and GLT-1 alanine mutants were expressed in HEK PS DKO at comparable levels to PS1wt and GLT-1 wt, respectively.

**Figure 1.**
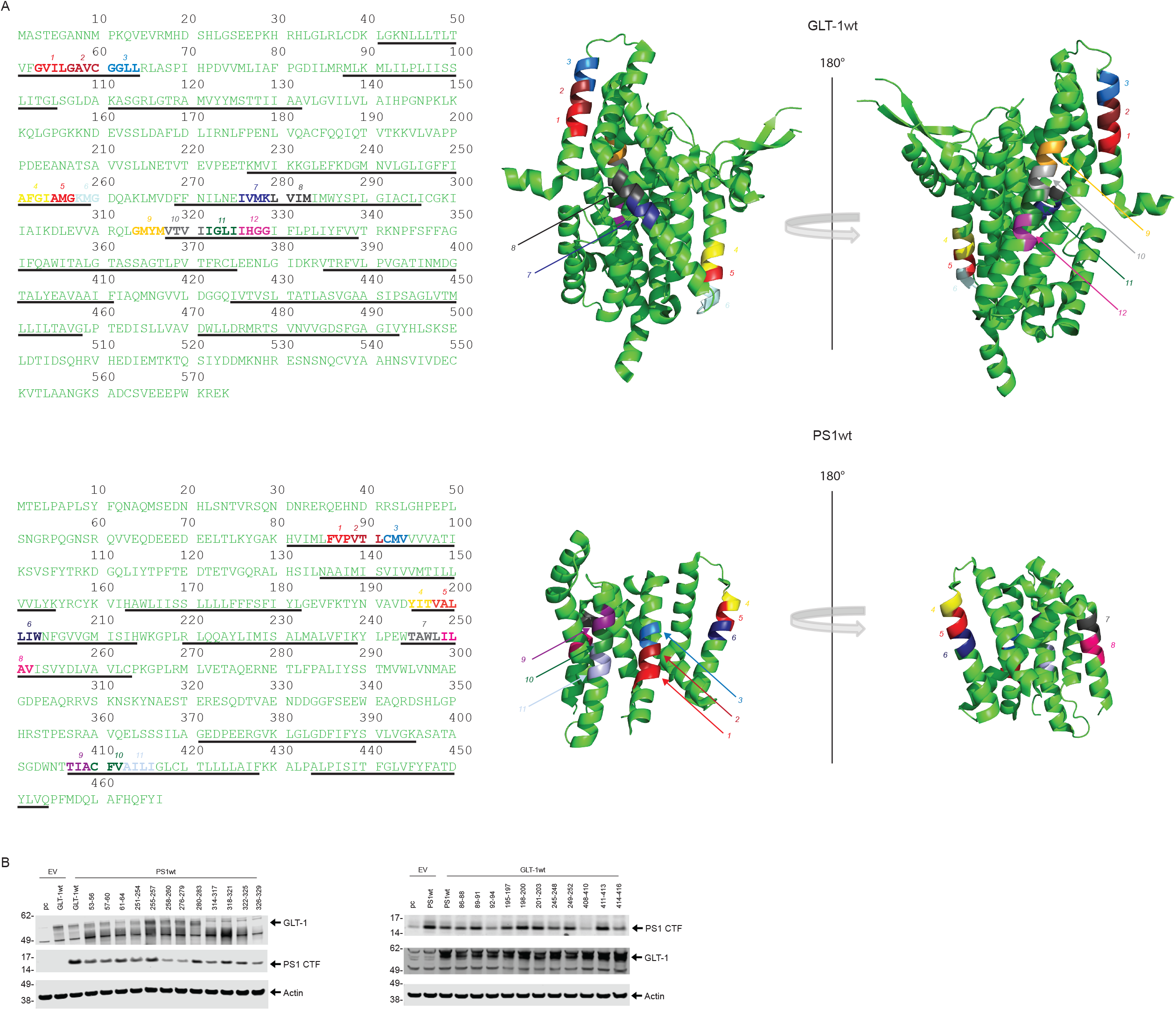
Experimental design for alanine scanning. mutagenesis to determine specific GLT-1 and PS1 amino acids representing potential interaction sites. A) Amino acid sequence of GLT-1 (top) and PS1 (bottom). Amino acids that can potentially interact with PS1 or GLT-1 are in bold and different colors; rest of the sequence is indicated in light green. Underlined amino acids correspond to transmembrane regions. Structure of GLT-1 and PS1 (adapted from PDB: 7XR4 and PDB: 5A53 respectively). Two representations with a rotation of 180°. B) Representative western blots showing the level of expression for wild type (wt) and alanine mutant GLT-1 (left) and PS1 (right) and in HEK PS DKO cells. The cells were transiently co-transfected with empty vector (EV) and wtPS1(left) or wtGLT-1 (right).

### Identification of the interaction site between PS1 and GLT-1

To test whether GLT-1/PS1 interaction is disrupted by PS1 and GLT-1 alanine mutations, we employed FRET-FLIM approach in intact CHO cells transiently co-transfected with either GLT- 1 alanine mutants and PS1 wt or PS1 alanine mutants and GLT-1 wt. The cells were immunostained with anti-PS1 and anti-GLT-1 antibodies, followed by AF488 and Cy3-labelled secondary antibodies, respectively. Anti-GLT-1 antibody was omitted in one well, which served as negative FRET control. Donor fluorophore (AF488) lifetimes were measured as an indicator of proximity between GLT-1 and PS1. Shortening of the AF488 lifetime signal increases as a result of FRET, suggestive of energy transfer between fluorophores and consistent with protein-protein interaction (Figure 2A-C and Fig. S1). We found that GLT-1 alanine mutants at position 251-254 and 276-279 had significantly decreased FRET efficiency (i.e., interaction) with PS1wt (Figure 2 B). Similarly, we observed a significantly reduced interaction with GLT-1 wt for PS1 alanine mutants at position 249-252 (Figure 2D). We ran Alphafold Multimer to create a model of GLT-1/PS1 interaction and interestingly, two of the top 5 models suggested a GLT-1/PS1 interface between GLT-1 TM5 and PS1 TM6, where our alanine scanning showed a statistically significant decrease in FRET efficiency upon mutation. The model presented shows GLT-1 residues 274 to 281 and PS1 residues 247 to 254 are engaged in the interface (Figure 2E).

**Figure 2.**
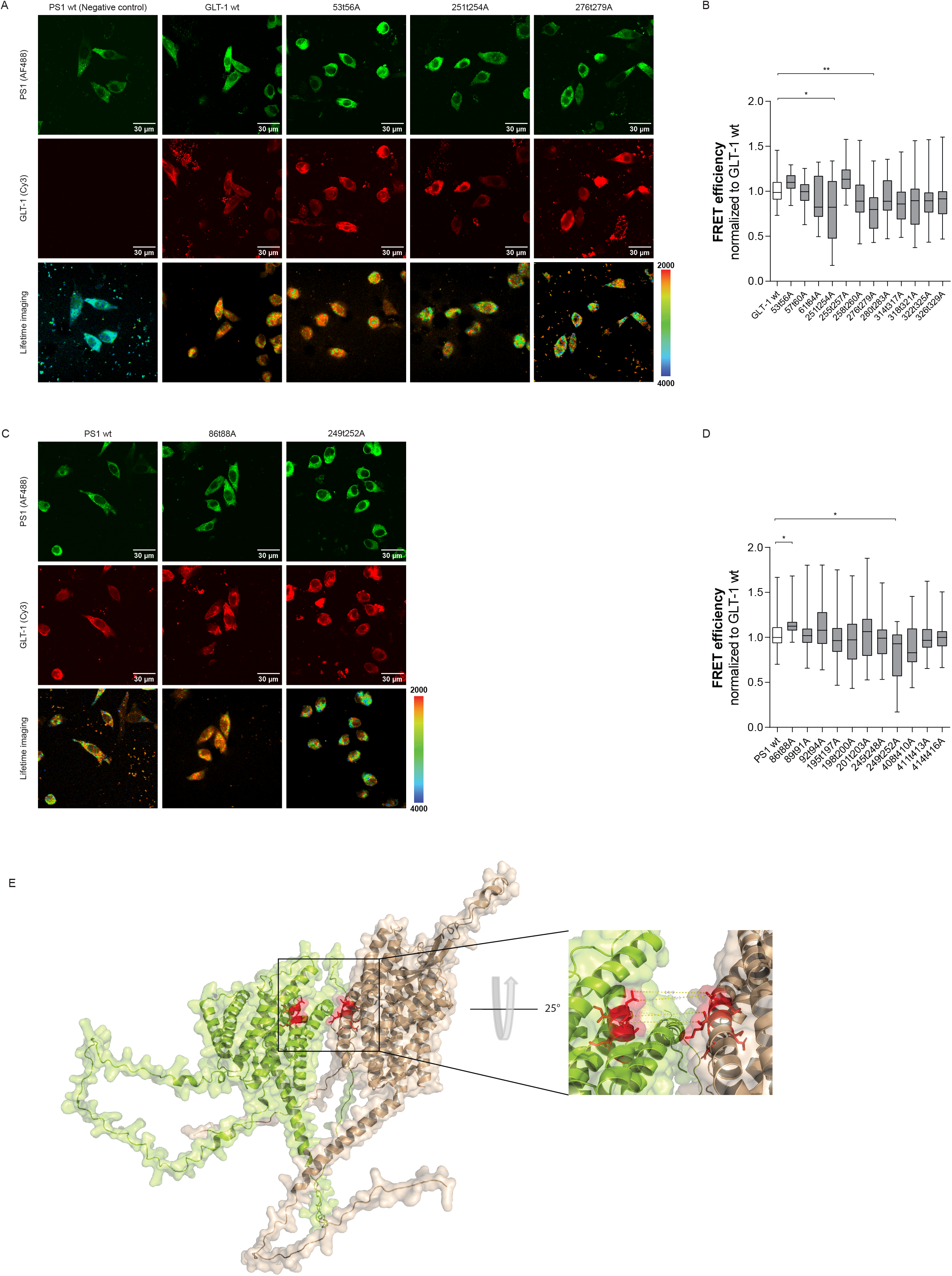
Identification of GLT-1/PS1 interaction sites by Fluorescence Lifetime Imaging Microscopy, FLIM. A) Confocal images show GLT-1 wt and GLT-1 alanine mutants’ immunoreactivity in CHO cells. The cells were immunostained with anti-PS1 and anti-GLT-1 antibodies, followed by AF488 and Cy3-labelled secondary antibodies, respectively. Color-coded FLIM images show AF488 PS1 donor fluorophore lifetimes, representing different proximity between PS1 and GLT-1 fluorophore. Colorimetric scale shows fluorescence lifetime in picoseconds. Longer AF488 lifetimes (blue-green pixels) represent diminished proximity, i.e., reduced interaction between PS1 and GLT-1. B) FLIM analysis of GLT-1/PS1 proximity (%E_FRET_) in CHO cells. Average %E_FRET_ in cells expressing wt GLT-1 (n=4 independent experiments, ∼15 cells per ROI), and GLT-1 alanine mutants (n=4, 10∼cells per ROI). Box and whiskers plot with min and max bars; Kruskal-Wallis ANOVAs with Dunn’s multiple comparison test. C) Confocal images show PS1 wt and PS1 alanine mutants’ immunoreactivity in CHO cells. The cells were immunostained with anti-PS1 and anti-GLT-1 antibodies, followed by AF488 and Cy3-labelled secondary antibodies, respectively. Color-coded FLIM images show AF488 PS1 donor fluorophore lifetimes, representing different proximity between fluorophores labeling PS1 and GLT-1. Colorimetric scale shows fluorescence lifetime in picoseconds. D) Average %E_FRET_ in cells expressing PS1wt as a control (n=4, 15∼ cells per ROI), and GLT-1 alanine mutants (n=4 independent experiments, 10∼cells per ROI). The results are expressed as FRET efficiency normalized to the corresponding wild-type protein. Box and whiskers plot with min and max bars, Kruskal-Wallis ANOVAs with Dunn’s multiple comparison test. E) AlphaFold multimer model of GLT-1 and PS1 interaction with a zoomed in view to the interaction. PS1 is in lime green, and GLT-1 is in wheat with the potential interaction sites in red.

### Design of the Cell Permeable Peptide

After identifying the interaction site between PS1 and GLT-1, we designed cell-permeable peptides (CPPs) that should block the interaction sites on both GLT-1 and PS1. CPPs are short positively charged peptides composed of basic residues (lysine or arginine) of about 20 to 50 amino acids which can cross the cellular plasma membrane (26). Our design used the HIV TAT domain that forms an alpha helix fused via a triple glycine linker (27) to PS1 or GLT-1 sequences identified previously as the interaction site. For creating the GLT-1 CPPs, we went ahead with the 276–279 aa GLT-1 mutant as it had more pronounced reduction in GLT-1/PS1 interaction (Figure 3A). To confirm the cell-penetrating capacity of the CPPs, we designed two other identical peptides fused with FAM tag (Fluorescein amidites) and assessed the uptake of CPPs by mouse primary neurons using confocal microscopy. A similar approach has been successfully used previously in our lab to disrupt the interaction between PS1 and Synaptotagmin 1 (28).

**Figure 3.**
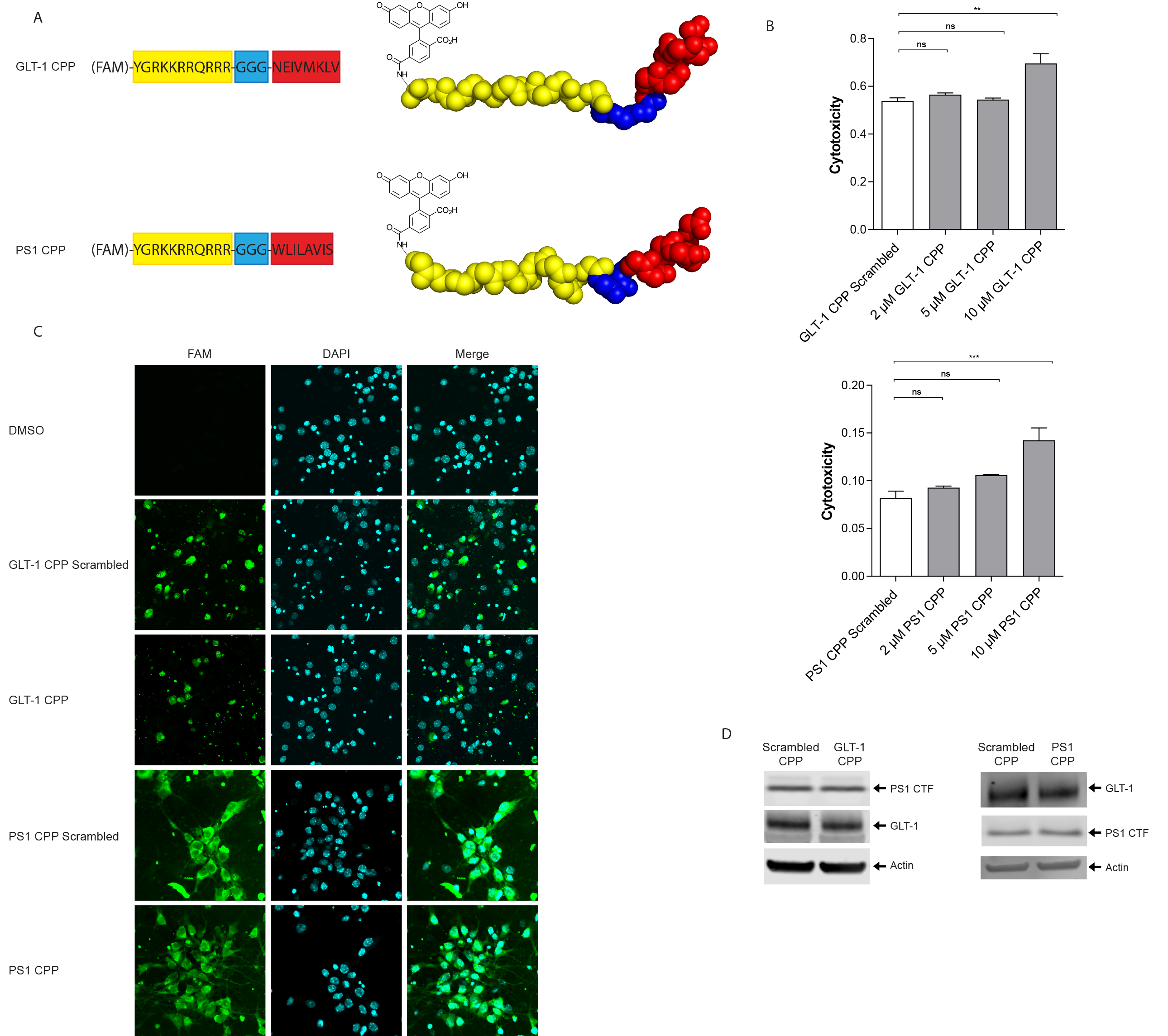
Experimental design for CPP. A) Schematic representation of the GLT-1 and PS1 cell-permeable peptides. The GLT-1 or PS1 fragment used for the design of the cell-permeable peptide aimed to inhibit the GLT-1/PS1 interaction is indicated in red. The scheme presents the fusion of the HIV1 TAT domain (in yellow) with the respective GLT-1 (top) and PS1 (bottom) fragment (in red) via GGG linker (in blue). For initial experiments, the peptides were conjugated with fluorescein (FAM) (in black) to visualize intracellular delivery. B) Cytotoxicity was measured in mouse primary neurons at DIV10 after a two-hour incubation at the respective concentrations. Cytotoxicity was determined using lactase dehydrogenase (LDH) activity assay. Toxicity of GLT-1, PS1, and scrambled peptides were determined at 2, 5, and 10 μM final concentrations. The data are presented as mean ± SEM, n =3 for 2 μM, n =6 for 5 μM and, n =3 for 10 μM. Statistical significance was determined using One-way ANOVA with Dunnett’s multiple comparison test. C) Representative images demonstrate successful delivery of 5 μM FAM-TAT-GLT-1, FAM-TAT-PS1, and FAM-TAT-scrambled peptides (all in green) to the primary neurons after a two-hour incubation. The cells were counterstained with DAPI (blue) to visualize the nuclei; scale bar 30 μm. D) Representative western blots showing GLT-1 and PS1 expression in neurons after 5 uM CPPs’ treatment. Membranes were probed for GLT-1, PS1, and actin.

The potential toxicity of these two CPPs and their respective scrambled (control) peptides was measured in primary mouse neurons at DIV10–12 after a two-hour treatment at 2, 5, and 10 μM final peptide concentrations. The percentage of cytotoxicity following the treatment was determined using the LDH activity assay. We did not notice significant toxicity for either CPP at 2 or 5 μM, while both CPPs at 10 μM showed some toxicity (statistically significant) (Figure 3 B). To look for the efficacy of these CPPs in downregulating the GLT-1/PS1 interaction, we chose 5 μM moving forward as this concentration was not toxic to the cells.

To check the CPPs’ cell-penetrating capacity, we treated neurons for two hours with 5 μM FAM tagged CPP. GLT-1, PS1, and scrambled CPPs were incubated with primary neurons for 2 hours; FAM-positive fluorescence was verified by confocal microscopy (Figure 3C).

Next, we assessed if these peptides can change PS1 or GLT-1 expression after internalization. Neurons were exposed to 5 μM GLT-1 CPP or PS1 CPP for two hours and then lysed.

Immunoblots were probed for GLT-1, PS1, and β-actin antibodies. No difference was observed in terms of relative GLT-1 or PS1 expression level (Figure 3D).

### CPPs block GLT-1/PS1 interaction

To determine whether PS1 and/or GLT-1 CPP would inhibit the GLT-1/PS1 interaction, we measured the FRET efficiency in intact neurons at DIV10–12; corresponding scrambled CPPs were used as controls. After two hours of treatment with 5 μM CPPs, neurons were fixed and probed with PS1 and GLT-1 primary antibodies, and Alexa fluor 488 and Cy3 secondary antibodies respectively (Figure 4A). FRET efficiency is indicative of short or long proximity between the fluorophores labeling PS1 and GLT-1. Scramble peptide-treated neurons were comparable to vehicle-treated neurons (Figure 4B). We observed a significant decrease in GLT-1/PS1 interaction in GLT-1 CPP and PS1 CPP treated neurons, compared to neurons treated with their respective scrambled CPPs. We illustrated CPPs binding with their specific targets (Figure 4C).

**Figure 4.**
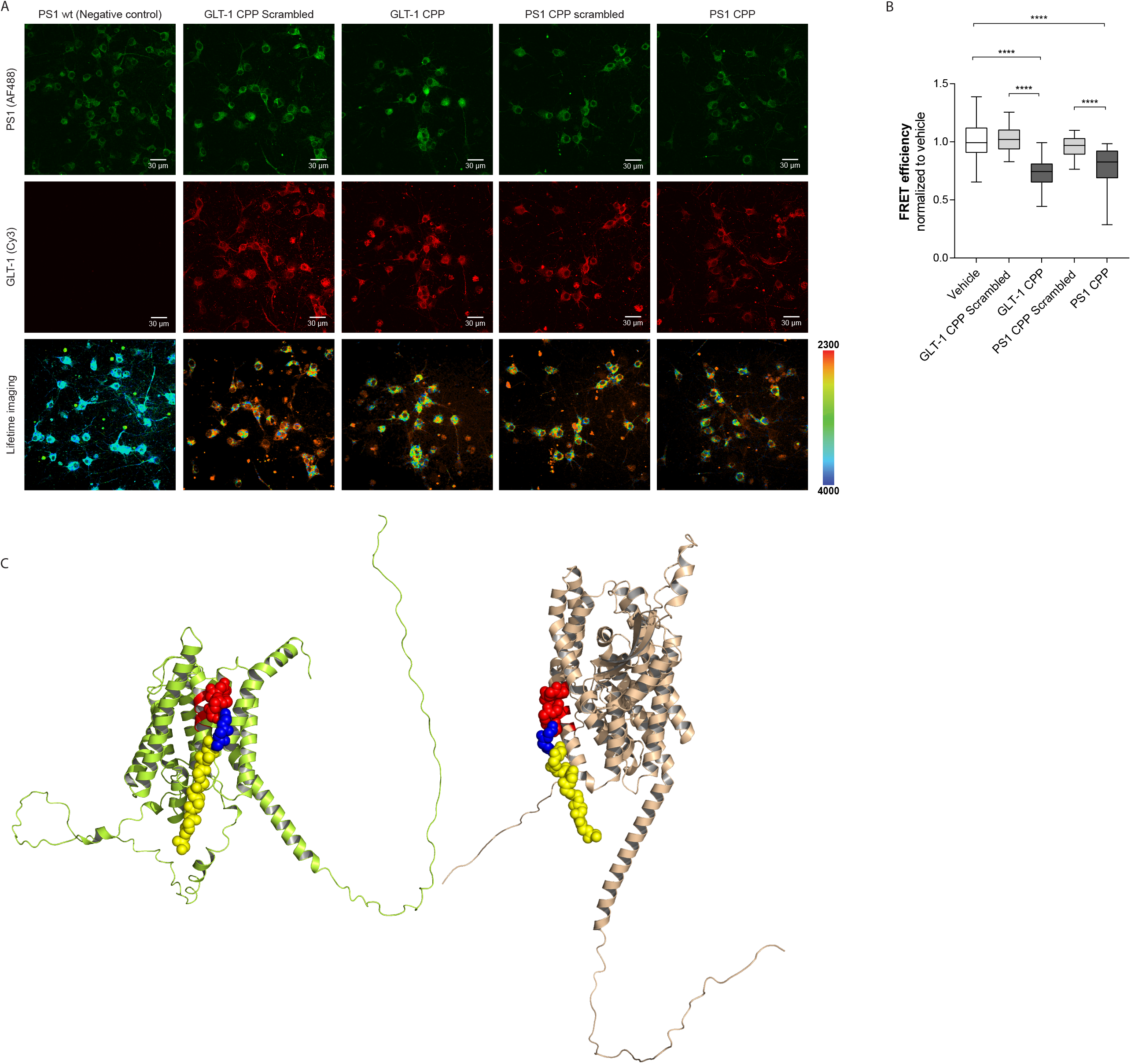
CPPs effects on PS1 and GLT-1 interaction by FLIM in primary neurons. A) GLT-1/PS1 proximity in neurons treated with 5 μM GLT-1 CPP, PS1 CPP, or corresponding scrambled peptides for two hours was determined by FLIM. The neurons were immunostained with anti-PS1 and anti-GLT-1 antibodies, followed by fluorescently conjugated secondary antibodies AF488 and Cy3, respectively. Confocal images show endogenous GLT-1 and PS1 immunoreactivity (top two rows). The pseudo-colored images (bottom row) represent the donor fluorophore lifetime in picoseconds. The orange-red pixels indicate shorter lifetimes reflecting closer proximity between the fluorescently labeled PS1 and GLT-1 B) The change in FRET efficiency was used to estimate relative change in the proximity between PS1 and GLT-1. The graph shows normalized FRET efficiency between GLT-1 and PS1. The data are presented as mean ± SEM, n = 3 independent experiments for GLT-1 and PS1 scrambled CPPs, n = 6 for GLT-1 CPP and PS1CPP. Statistical significance was determined using Kruskal-Wallis ANOVAs with Dunn’s multiple comparison test; **** = p <0.0001 and ns = non-significant. C) 3D models of GLT-1 CPP and PS1 CPP binding with PS1 and GLT-1 respectively. Models used AlphaFold predictions and have been assembled on Pymol. PS1 is in lime green, and GLT-1 is in wheat. CPPs are in yellow, blue, and red representing TAT domain, linker, and interacting sequence, respectively.

## Discussion

We recently found that PS1 and GLT-1 interact *in vivo* in the mouse brain, and this interaction occurs in both astrocytes and neurons (19). In the current study, we unraveled the interaction sites between PS1 and GLT-1 and designed CPPs to block this interaction in primary mouse neurons without altering the proteins’ expression levels. We found that human GLT-1 residues 251-254 and 276-279 (Figure 2B) interact with human PS1 residues 249-252 (Figure 2D).

The PS1 structure was resolved in 2015 as a γ-secretase complex component (29), which we used to build interaction probability models. Our predictions about GLT-1/EAAT2 structure were initially based on GLT-1 analogs such as the sodium/aspartate symporter from *Pyrococcus horikoshii* (Glt_Ph_) (30, 31). The structure of GLT-1 has been recently identified using cryo-EM (32, 33), corroborating our model with predicated sites where PS1 could be physically interacting with GLT-1 and vice versa. We mainly focused our alanine scanning on accessible residues in folded PS1 and GLT-1 proteins and the juxta-transmembrane region with hydrophobic domains and nonpolar residues. These properties lead to electrostatic interactions driving folding and protein stability (34). Based on our study, GLT-1 interaction sites with PS1wt are at position 251 to 254 and 276 to 279 which correspond to GLT-1 transmembrane (TM) 4 and TM5, respectively. According to Reyes study on Glt_Ph_ (35), TM2, TM4, and TM5 are responsible for all inter-subunit contacts. Furthermore, these domains form a highly accessible scaffold domain (32, 33). TM5 in position Trp286 present cholesterol binding domain. Plausibly, this region facilitates contact with larger proteins such as PS1. Mutation of GLT-1 Trp286 to alanine showed that glutamate uptake was reduced (32) suggesting that Trp286 mutation has negative effects on the transport activity in the plasma membrane environment.

Since Trp286 is critical for GLT-1 function, we can hypothesize PS1 binding at nearby amino acids 276–279 may modulate GLT-1-mediated transport activity due to steric hindrance, changing the flexibility of the GLT-1 TM5 alpha helix.

Conversely, PS1 residues 249-252 in TM6 interact with GLT-1. PS1 transmembrane domain 6 is the most flexible domain and can adopt several conformations (36) which may facilitate its interaction with GLT-1. The PS1/γ-secretase autoproteolytic cleavage sites are at positions 257 and 385, located in TM6 and TM7 respectively. These endoproteolysis sites are critical for presenilin maturation and function, with the exception of PS1ΔE9 fAD mutants that are enzymatically active in the absence of endoproteolysis (37). In our previous study, we reported a successful pull-down of PS1-CTF and PS1 full length with an anti-GLT-1 antibody (19). This indicates that GLT-1 may interact with both PS1 forms, mature and immature, regardless of endoproteolysis. Furthermore, the interaction site is also very close to a mutation hotspot located in exon 8 such as the Paisa mutation (E280A) related to epilepsy (38), demonstrating the importance of this interaction in AD, particularly fAD. We can extrapolate that PS1 mutations impair the interaction with GLT-1 leading to increased epileptiform activity observed in AD patients (6-8) (39).

However, we cannot deny the possibility that the interaction may occur at other positions involving polar residues as well. A more robust approach such as cryo-EM could provide clearer information about the precise interaction site(s) between the two proteins as well as GLT-1/PS1 folding when they interact with each other. Such information could help solidify our CPP design to affect this interaction positively or negatively in an AD context. We ran AlphaFold2 to model both proteins while interacting with each other, and two of the models produced the same interface as our alanine scanning results (Figure 4C), cross validating the location of the interaction site we identified.

Successful identification of the PS1 and GLT-1 binding sites allowed us to create a tool to modulate GLT-1/PS1 interaction using CPPs. We show that TAT-CPPs based on the identified interaction sites penetrated the plasma membrane targeting PS1 and GLT-1 interaction without affecting the proteins’ expression levels (Figure 3D). The blood brain barrier crossing property of TAT-CPPs is widely used for brain delivery of drugs (40), and can be harnessed for AD therapeutics targeting PS1/GLT-1.

In summary, pinpointing the GLT-1/PS1 interaction sites is crucial for unraveling the significance of this recently uncovered interaction between PS1 and GLT-1, and provides a first step to solve the connection between aberrant glutamate transport and amyloid pathology in AD. Alleviating glutamate homeostasis impairments and modulating Aβ deposition, both early events preceding clinical manifestation of AD, may represent a new therapeutic avenue for AD. Thus, developing a tool, such as PS1/GLT-1 CPP, to modulate the interaction between GLT-1 and PS1 in their native environment within cells is a crucial step for uncovering the significance of this interaction, providing therapeutic potential for AD.

## Acknowledgements

This work was supported by NIH AG15379 (OB and RET), AG44486 (OB), R01 AG055784 (CZ), and Cure Alzheimer’s Fund (RET). Thank you to Dr Calina Glynn for proofreading of the manuscript.

## Author contributions

F.P. and O.B. conceived and designed the experiments, F.P., P.S. and S.M. collected and analyzed the data, M.M. provided helpful discussion; F.P., P.S. and O.B. wrote the paper.

## Figures legends

**Figure S1. FLIM analysis of the PS1/GLT-1 interaction.** The cells were immunostained with anti-PS1 and anti-GLT-1 antibodies, followed by AF488 and Cy3-labelled secondary antibodies, respectively. Color-coded FLIM images show AF488 PS1 donor fluorophore lifetimes, representing different proximity between PS1 and GLT-1 fluorophore. Colorimetric scale shows fluorescence lifetime in picoseconds. Longer AF488 lifetimes (blue-green pixels) represent diminished proximity, i.e., reduced interaction between PS1 and GLT-1. Pseudo-colored FLIM images show interaction between PS1 wt and GLT-1 wt/GLT-1 alanine mutants (A), and between GLT-1 wt and PS1 wt/PS1 alanine mutants (B).

