## Supplementary figures and images for "Identification of PS1/gamma-secretase and glutamate transporter GLT-1 interaction sites"

### Supplemental Figure 1

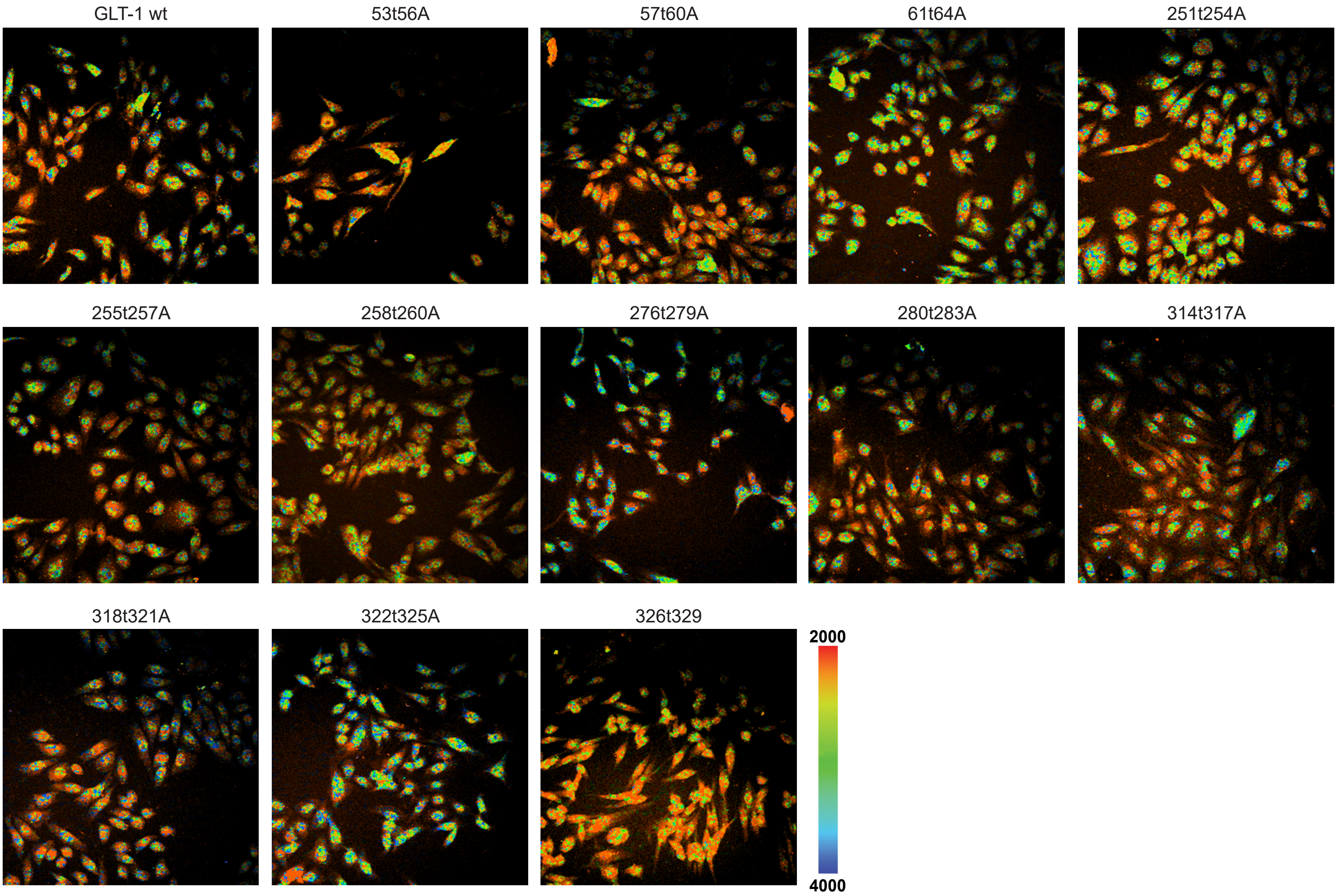

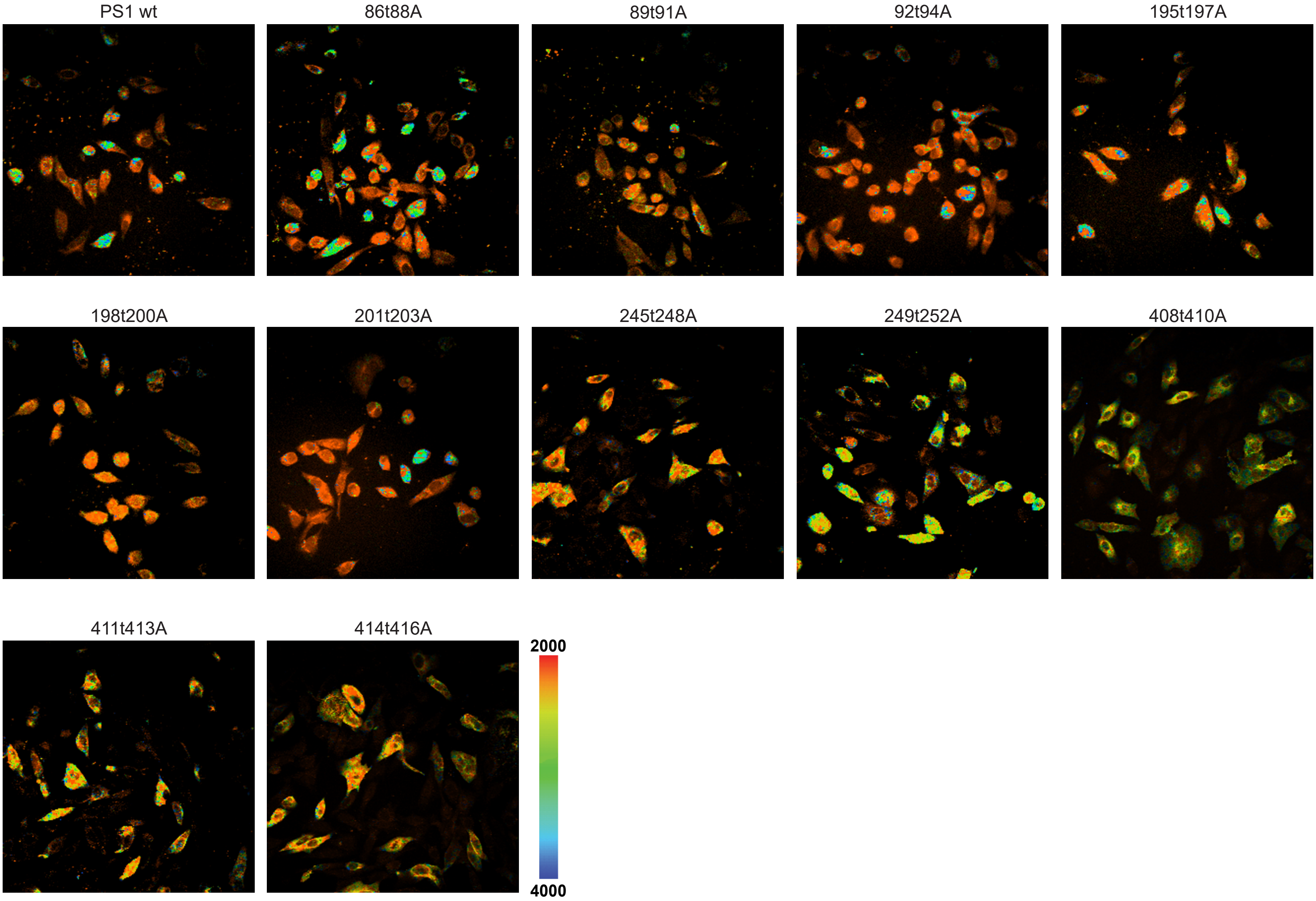
